# Data Gap or Biodiversity Gap? Evaluating apparent spatial biases in community science observations of Odonata in the east-central United States

**DOI:** 10.1101/2022.11.29.518107

**Authors:** Christian M. Bullion, Christie A. Bahlai

## Abstract

Odonates (dragonflies and damselflies) have become popular study organisms for insect-based climate studies, due to the taxon’s strong sensitivity to environmental conditions, and an enthusiastic following by community scientists due to their charismatic appearance and size. Where formal records of this taxon can be limited, public efforts have provided nearly 1,500,000 open-sourced odonate records through online databases, making real-time spatio-temporal monitoring more feasible. While these databases can be extensive, concerns regarding these public endeavors have arisen from a variety of sources: records may be biased by human factors (ex: density, technological access) which may cause erroneous interpretations. Indeed, records of odonates in the east-central US documented in the popular database iNaturalist bear striking patterns corresponding to political boundaries and other human activities. We conducted a ‘ground-truthing’ study to examine these patterns in an area where community science reports indicated variable abundance, richness, and diversity which appeared to be linked to observation biases. Our observations were largely consistent with patterns recorded by community scientists, suggesting these databases were indeed capturing representative biological trends and raising further questions about environmental drivers in the observed data gaps.

## Introduction

Community science initiatives have been crucial for understanding changes in biodiversity, distribution, and phenology, due to their potential to generate large volumes of data and cover broad geographical areas (Fraisl et al., 2022). The possible benefits of community science have been well-documented and could represent a viable alternative to data acquisition for projects where scarce financial or logistical resources prevent traditional, multi-visit sampling (Lauret et al., 2021). For example, in one study, youth volunteers were found to have observed proportionally higher numbers of mollusks, arachnids, and insects than the average iNaturalist user (Aristeidou et al., 2021). While in the context of water quality, community science was able to produce consistent measurements and statistically validated data pertaining to groundwater and rainfall (Walker et al., 2016). The theoretical benefits of community science go on, with broader scientific literacy, low/no-cost data sources, and increased monitoring of underserved ecosystems being cited among many potential advantages in a large review of community science (Conrad & Hilchey, 2011).

Community science data may also represent a considerable boon to academic biodiversity science as a data source for species where formal data collection is rare or incomplete, such as taxa not considered to have economic importance. For example, odonates (dragonflies and damselflies) have fostered a large and widespread hobbyist following that has provided nearly 1,500,00 open-sourced odonate records worldwide through public databases like Odonata Central and iNaturalist— entries that could conceivably lay the groundwork for numerous insect ecological studies. The phenology and spatial distribution of the odonates is tied closely to environmental cues and conditions, primarily temperature and photoperiod, even leading to them being referred to as ‘living barometers’ (Hassall, 2015). However, climate-driven disruptions to the delicate ‘when’ and ‘where’ of odonates could have severe implications for these insects (Zarnetske et al., 2012). For example, range-shifting (as a species’ population shifts to areas with more favorable environments) and changes in phenology (the timing of life-history events) have both been well-documented climate-linked responses in odonates (Hassall & Thompson, 2008, 2010; Hickling et al., 2005; May et al., 2017; Winder & Schindler, 2004). Community science records have already been used, with great success, in the long-term monitoring of Californian odonates (Rapacciuolo et al., 2017a) and to identify breeding occurrences in Oklahoma odonates (Patten et al., 2019).

However, understanding these fundamental biodiversity trends requires data that is unbiased in time and space, and community science has not gone uncriticized in these regards (Catlin-Groves, 2012; Lukyanenko et al., 2016). Most concerningly, community science initiatives risk reflecting biases and interests held by their participants. In one study, areas of public concern were suggested to have been oversampled by community participants (Jollymore et al., 2017). Similarly, other studies have criticized user-generated geographical data, questioning their reliability, quality, and overall value (Flanagin & Metzger, 2008). However, these issues may not be easily addressed, as an increasing focus on data quality could come at the cost of widespread accessibility (Parsons et al., 2011).

Biases arising from infrastructure and human population density are of particular note. For example, in comparison to their observed biological richness, agricultural areas were vastly oversampled (Geldmann et al., 2016). Similarly, volunteer sampling can often be affected by a ‘cottage effect’, being more likely to sample easily accessible locations, such as those near roads or population centers (Millar et al., 2019). These patterns of interaction can have profound implications: research involving bird species have suggested that biases in community science data may produce less accurate models for areas with distinct environmental characteristics and low observation rates (Johnston et al., 2020). In their most extreme, these low-density areas may form complete data gaps, which interfere with our understanding of assemblages and species distributions, especially in areas that are highly vulnerable to diversity loss and highly understudied (Archer et al., 2014).

Indeed, our research group encountered a striking example of what appeared to be a geographical bias in reporting frequency for community science records when initiating a study utilizing community science records to examine recent changes in odonate communities in the east-central United States (Figure 1). In a prominent show of extreme sampling, the state of Ohio showed high-density reports across nearly its entire region, with state borders clearly visible on observation density heat maps. This pattern is driven, at least in part, by the popular Ohio Odonate Survey, a large-scale community science initiative that first ran from 1991 to 2001 and was re-initiated in 2017. In total, this initiative has reported over 125,000 odonate observations to iNaturalist (INaturalist, 2021; Ohio Dragonfly Survey, 2021).

**Figure 1.**
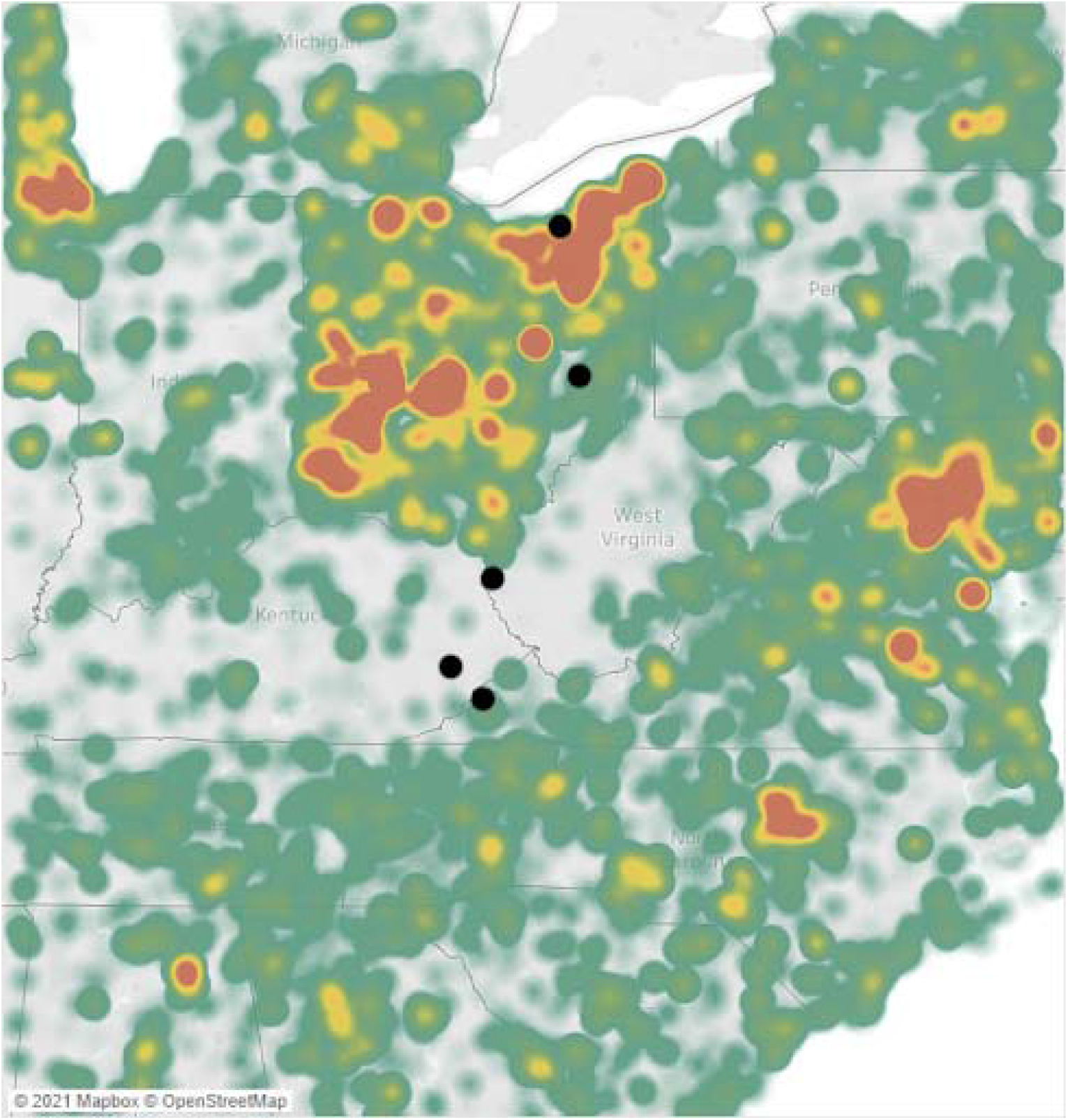
Density map of odonate community science observations, centered on the eastern United States. Green areas indicate low areas of reporting density while red indicates area of high reporting density. Survey sites are designated with black points.

In striking contrast, a large area of the central Appalachian region of the United States to the direct south, centered around West Virginia, is seemingly extremely underrepresented in reports for, not only odonates, but a variety of taxa in community science databases (INaturalist, 2021). This region is characterized by high elevation, temperate forests, and low human population densities, but has also experienced lower household incomes and higher poverty rates than surrounding regions in recent years (The Census Bureau, 2021). While this region neighbors the National Radio Quiet Zone (NRQZ), a federally designated region of West Virginia in which radio transmissions are heavily restricted, this NRQZ does not appear to contribute to the observed data gap, potentially due to a heightened focus on scientific research within that area. Meanwhile, this region of underreporting is echoed in the published ranges of several odonate species, recent community observations in that region have revealed the presence of several species considered to be at the extreme edge of, or outside of, their published ranges.

As such, we hypothesize that the observed data gap in central Appalachia is an artifact of human data collection patterns, particularly one arising from the inherent challenges facing observation-based community science initiatives in prominently rural areas: essentially, we predict that odonate biodiversity and abundance in these regions are being under-reported by community scientists compared to areas with higher populations and access to more economic resources. If so, these data collection artifacts could be shaping currently accepted species distributions through the strength of observation efforts that characterize them. Thus, we predict that between-site community trends will differ by human population density in the citizen science data but will be more equally distributed across sites in controlled expert surveys. Therefore, in this study, we set out to evaluate the reliability of community science surveys in documenting odonate diversity relative to expert sampling. As such, this study documents a ‘ground truthing’ study where expert sampling was paired with community science records on a north-south transect, to examine any discrepancies in abundance, diversity, and community composition produced by the two data collection methods.

## Materials & Methods

### Study Area

To evaluate patterns observed in community science records for this system, we conducted a ground-truthing survey based on comparing systematic ‘expert’ observations with publicly reported community science observation data from two major odonate open data sources, iNaturalist and Odonata Central. A north-south transect, spanning from northern Ohio to the central Appalachians and centering on the observed data gap, provided the foundation for our ground-truthing dataset and incorporated longitudinal and elevation aspects.

Sampling was completed in five counties along this transect-Cuyahoga County (OH), Guernsey County (OH), Wayne County (WV), Knott County (KY), and Wise County (VA)-chosen to be approximately equally spaced, with bodies of water in naturalized areas that were reasonably accessible from roads or campgrounds (Figure 1; Table 1). These human-accessible sites were selected as those which would reasonably represent areas where community scientist reporting would most likely occur for these counties and had physical attributes associated with ‘good’ habitat for many odonate species (open bodies of water with vegetated shores).

**Table 1.** Geographic information for the five sampling locations selected for field-truthing, arranged from north to south.

| County Code | County Name | Latitude (°N) | Avg. Elevation (m) |
| --- | --- | --- | --- |
| A | Cuyahoga (OH) | 41.4 | 174 |
| B | Guernsey (OH) | 40.1 | 311 |
| C | Wayne (WV) | 38.2 | 296 |
| D | Knott (KY) | 37.3 | 382 |
| E | Wise (VA) | 37.0 | 687 |

The transect was sampled once monthly during June, July, and August, representing the peak odonate flight season for this region during 2019. For each site, data were collected from three vantage points along the lake shoreline deemed reasonably accessible to hobbyists by foot. For each vantage point at a site, surveying was conducted for ten consecutive minutes, once during the peak of the day (11:00 - 12:00 EST) and again during the evening of the same day (17:00 – 18:00 EST), to increase the likelihood of surveying both diurnal and crepuscular populations.

### Odonate Sampling

For each sampling period, the number of odonates visible from the selected vantage point, facing towards the lake, was counted per species. Identification of individuals was primarily done in-hand via netting, using the Dragonflies and Damselflies of the East field guide for reference (Paulson, 2011), while identification of visually distinct species was done through observation only. Sampling was conducted by two personnel, one as the observer and another as the recorder, and was consistent across sites.

Community science data was represented by a combined Global Biodiversity Information Facility dataset, containing reports originating from iNaturalist and Odonata Central (GBIF.org, 2021). Dataset consisted of Odonate abundance data during the peak adult flight season (June, July, August) for the years 2014 – 2021 in the focal region. Data were subsetted to create ‘observation units’ corresponding to our systematic surveys: records were aggregated by county of record and month of capture. Because variable reporting in some areas created zero-biased data incompatible with community analysis, we combined data from multiple years to represent a ‘typical’ community that could be observed in a given place, at a given time of year.

### Quantification and Statistical Analysis

All statistical analyses and plotting were conducted using R software (R Core Team, 2021), using the following packages: *rgbif, plyr, ggplot2, vegan, MASS, broom, and BiodiversityR* (Chamberlain & Boettiger, 2017; Kindt & Coe, 2005; Oksanen et al., 2022; Robinson et al., 2022; Venables & Ripley, 2002; Wickham, 2011, 2016, p. 2).

Community observation data for the focal counties was obtained from iNaturalist and Odonata Central via GBIF alongside the observations collected from the transect samples detailed above. Subsets of these datasets were created to include species identity, county, and date for each observation. For convenience and ease of understanding, specific county names were omitted and replaced with letter coding, labeling the sampled counties as A, B, C, D, and E, in order from north to south (Figure 1, Table 1). For each dataset, count data were aggregated using the *plyr* package to form counts of each species per county per source. Before merging, each dataset was amended to include its source, differentiating between community observations and transect observations.

To evaluate species richness by method and source, Shannon diversity for each metric was calculated using the *vegan* package from the row sums of the merged dataset, with corresponding plots generated from the results. Further aggregation of the data allowed for comparisons of species counts by county per source per month. Generalized linear models (GLM) for species richness by source, county, and month were constructed using the *MASS* package from this aggregated data and included both a model with an interaction between source and county and one without. Model selection, through comparison of model AIC scores, was then used to evaluate the relationship of species richness counts between source and county; a model with a better fit when the interaction was included suggests that there are geographical patterns in the tested metric between observational methods. Likewise, a better-fitting model without an interaction effect would suggest that the same patterns are held between sites. This process was then repeated to evaluate odonate abundance from the same data set, with all associated plots generated using the *ggplot2* package.

To examine differences in community composition among sites, sampling periods, and sampling approaches, non-metric multidimensional scaling (NMDS) using Bray-Curtis dissimilarity metrics was conducted using the *vegan* package. Lastly, analyses of similarities (ANOSIM) and homogeneity of variances were also conducted using the *vegan* package.

## Results

This study included 1,573 observations, with 381 originating from structured sampling efforts and 1,192 originating from community science efforts. The blue dasher, *Pachydiplax longipennis*, was the most commonly recorded species of the 27 observed during structured efforts (13.9% of observations), while the ebony jewelwing (*Calopteryx maculate)* was the most commonly recorded species of the 82 observed via community science sources (7.9% of observations).

Combined observed odonate species richness was highly variable between sites, ranging from 11 to 70, with higher reported diversity at the extreme ends of the transect (Figure 2). Community sampling efforts reported the highest richness, at 82 species, compared to the 27 species reported through structured sampling for the same timeframe. Shannon diversity values reflected this, with the highest proportional richness in the transect extremes and through community sampling (Figure 3, 4). Both sampling methods neared, but did not reach, saturation for species richness, as illustrated by the species accumulation curves (Figure 5).

**Figure 2.**
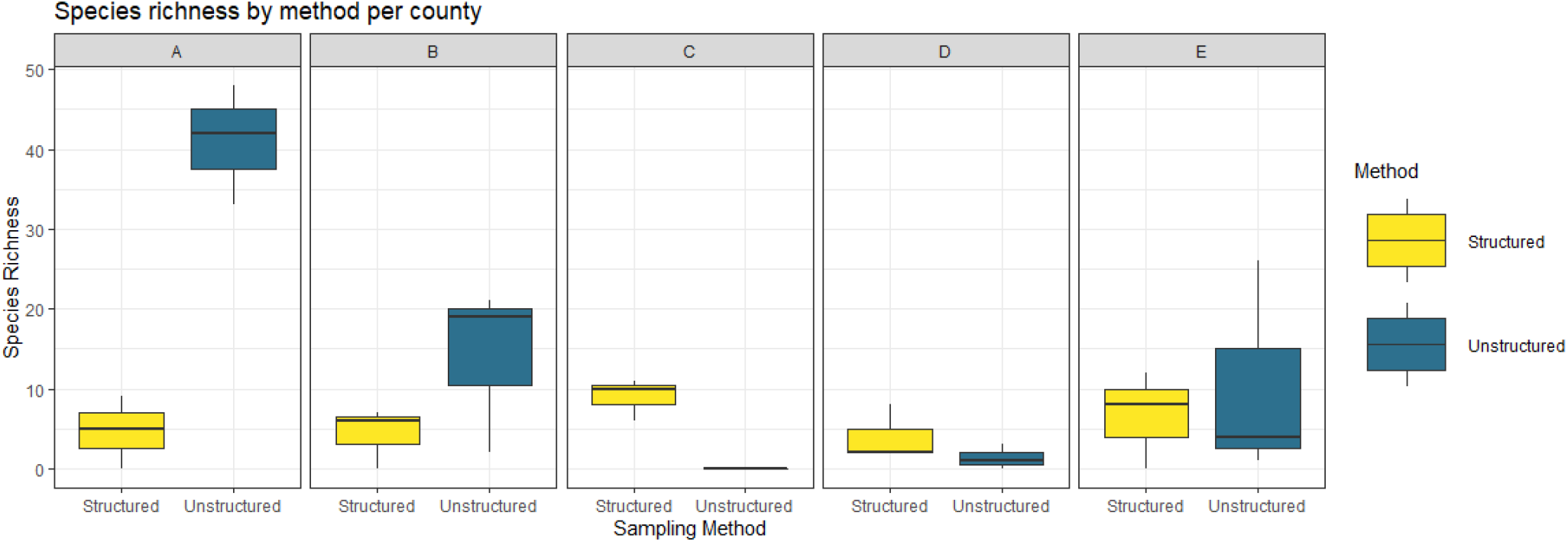
Boxplot comparing observed species richness across counties for each method.

**Figure 3.**
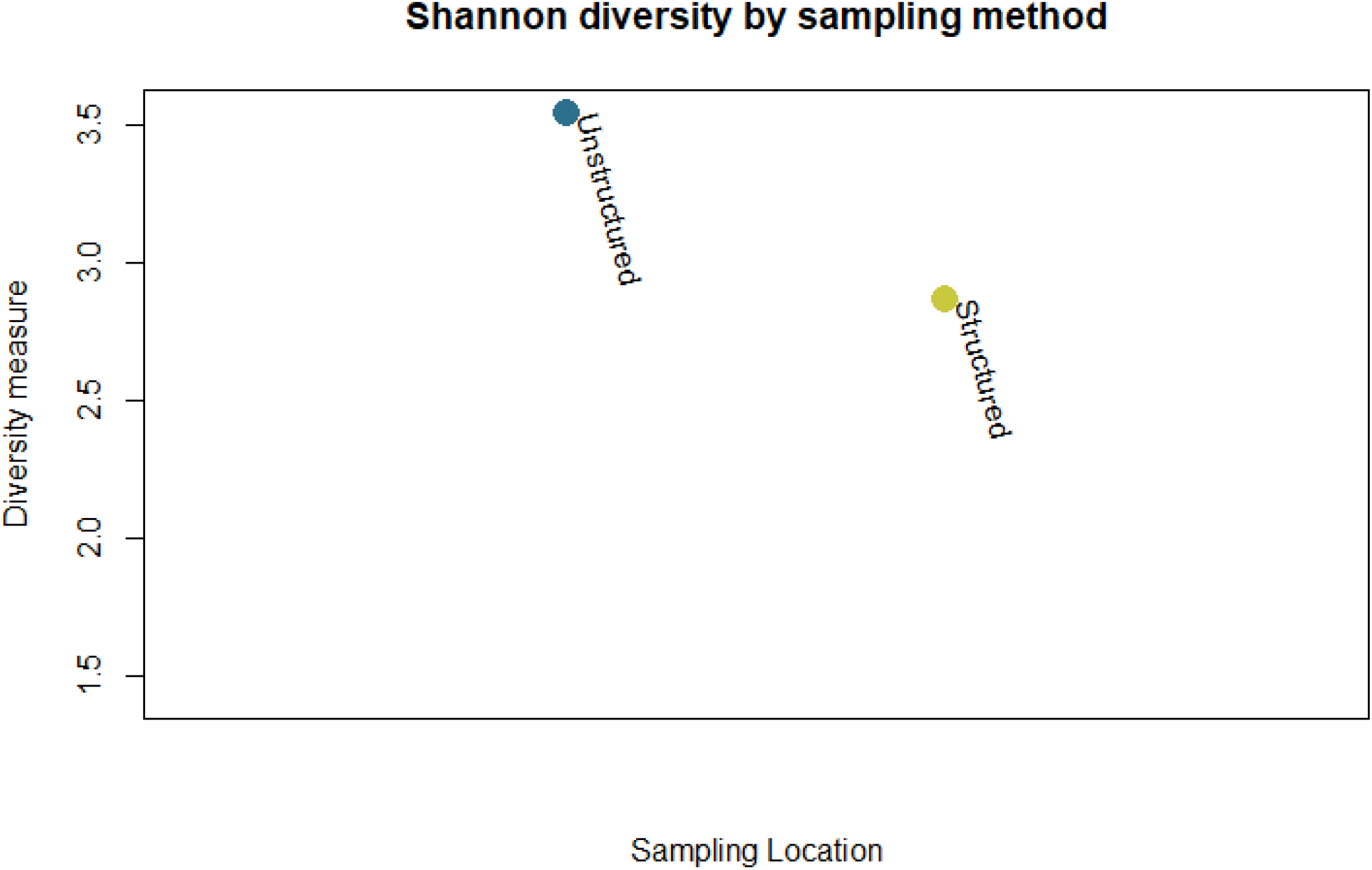
Shannon diversity by sampling method.

**Figure 4.**
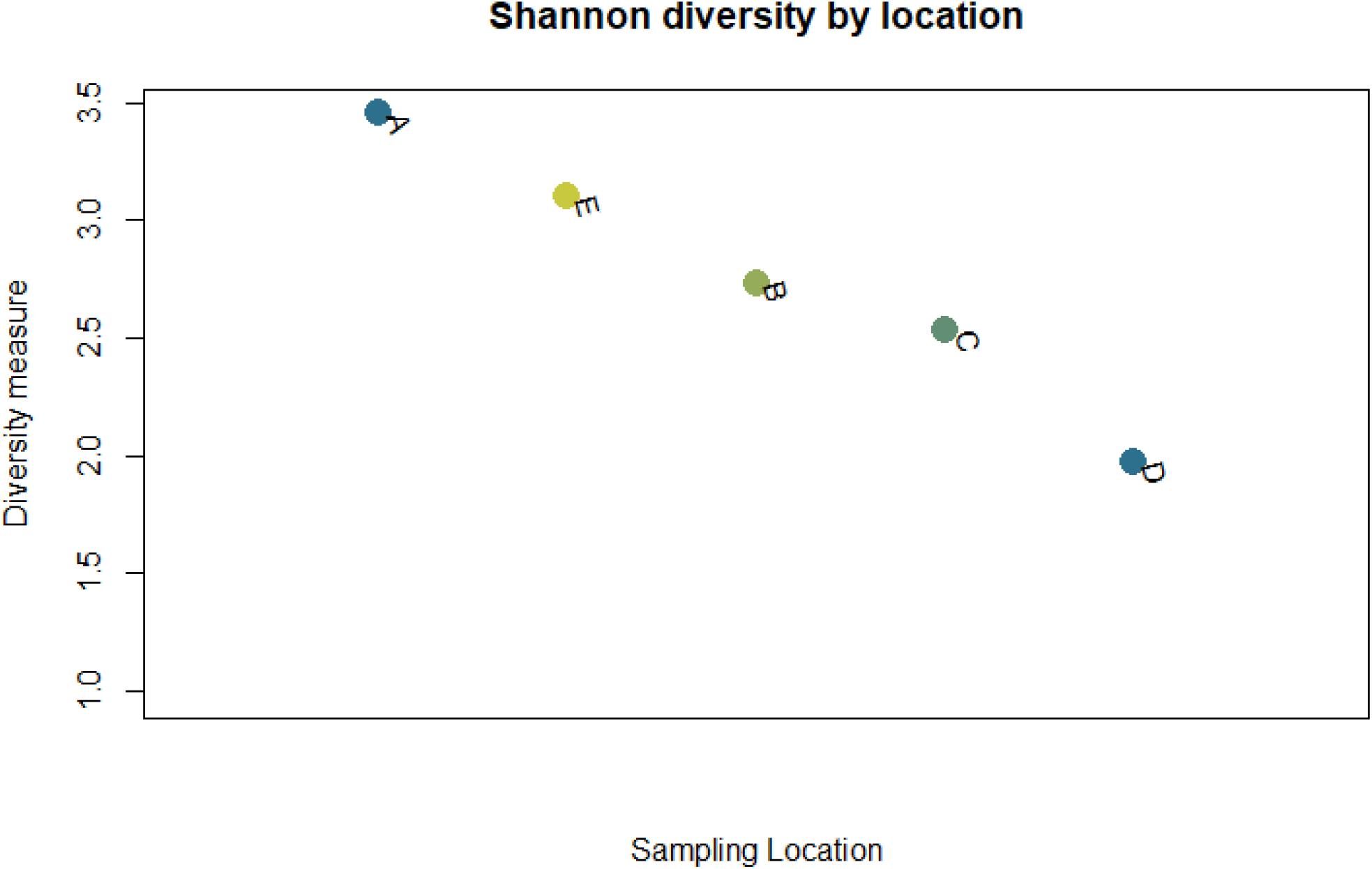
Shannon diversity by sampling location.

**Figure 5.**
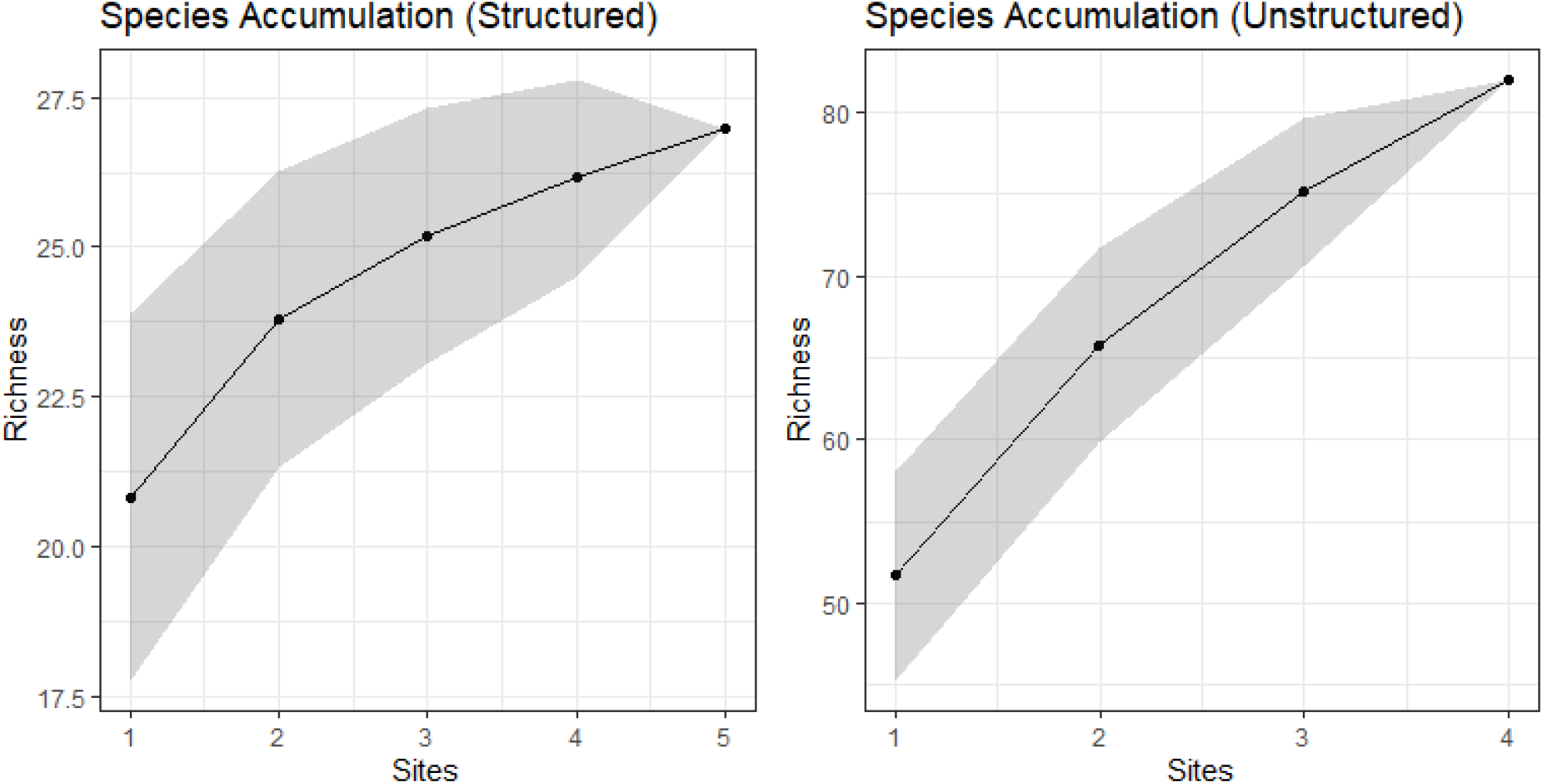
Species accumulation curves for both structured and unstructured sampling methods.

For generalized linear models of odonate species richness, the model with an interaction effect (AIC: 186) between method and sampling site performed better than the model without (AIC: 201), suggesting that there are different patterns in reporting behavior between sites per method. Structured sampling methods performed relatively consistently between sites while unstructured sampling methods had more variation, with each extreme end of the transect having more reported species richness than the intermediate data gap region. Additionally, a two-sided Student’s t-test suggested no difference between richness observations by method (p = 0.110).

Odonate abundance, measured as the total number of observations per species, was evaluated for each site and sampling source (Figure 6). Like the results obtained in the richness analysis, a generalized linear model that included an interaction between method and sampling site (AIC: 252) performed better than the model without (AIC: 262), suggesting that there were different patterns in overall abundance between sites per method. Student’s t-test found no difference in abundance by method (p = 0.254).

**Figure 6.**
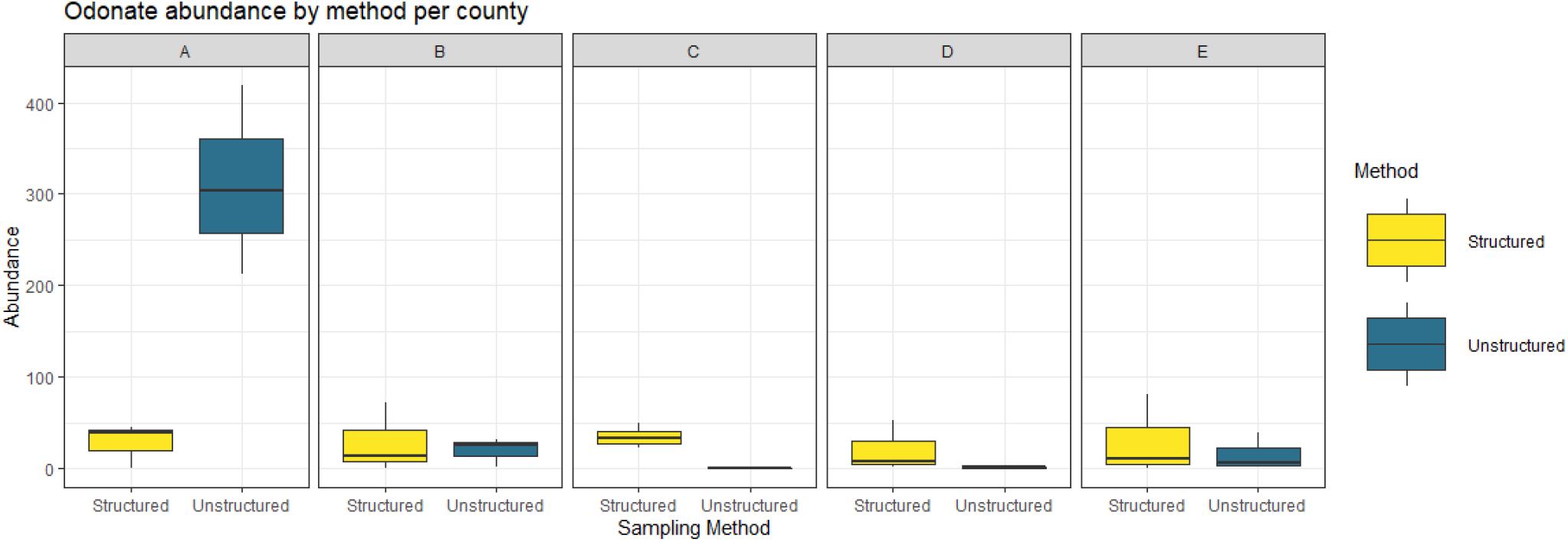
Boxplot comparing observed species abundance across counties for each method.

Community science and structured in-person sampling efforts captured similar odonate species composition at the same sampling sites (ANOSIM, p = 0.07). Likewise, beta dispersion analysis revealed that community science observations had more variable composition than structured sampling efforts; however, analysis of variation did not find these to be significantly different (p = 0.466).

## Discussion

For the focal counties and time periods, we found that unstructured community science and structured ‘expert’ sampling performed comparably well in reporting Odonate biodiversity. Both sampling methods reported population composition similarly, though structured sampling showed less variability. In the unstructured data set, reported species richness and abundance varied across the transect route, with extreme ends showing higher measurements than locations intermediate. However, significant discrepancies in the reporting patterns across the transect route for both methods suggest that the drivers behind reported biodiversity could extend beyond strictly anthropogenic forces.

The observed data gap was most apparent in the unstructured data set, with structured efforts reporting a more consistent, but less overall, diversity between sites. In contrast, unstructured efforts outperformed structured efforts in locations with more overall reports. This was particularly true for Ohio sites compared to non-Ohio sites, likely as a result of the efforts of the Ohio Odonate Survey, whose state-wide efforts have contributed over 150,000 open-sourced records over the last two decades.

Our findings agreed with numerous studies supporting the viability of community science in conservation (Lauret et al., 2021; McKinley et al., 2017; Walker et al., 2016) and Odonata biodiversity research (Patten et al., 2019; Rapacciuolo et al., 2017b). However, while inconclusive and of limited spatiotemporal scale, our observations of the data gap region mirror concerns about sampling biases and decision-making in community science that have also been the focus of many other studies (Archer et al., 2014; Bowler et al., 2022; Johnston et al., 2020; Millar et al., 2019; Ruete, 2015).

While community science records covered several years of flight seasons, field truthing efforts were only able to cover the peak season of 2019, as travel restrictions arising from the COVID-19 pandemic prevented repeated field sampling in the following years. Once-per-month field truthing allowed us to cover a much larger distance but introduced a risk of underreporting highly migratory species, like the green darner (*Anax junius)*, whose swarms tend to attract large numbers of community science reports nationwide. While this spatio-temporal snapshot is not broad enough to speak for the applications of community science as a whole, it has provided an interesting case study for an understudied region, upon which future studies can be built.

Lastly, we acknowledge that community science data is limited by the common usage patterns of their platforms. In particular, it is likely that the abundance reported by community science is much higher than actual values, as many community scientists report based on species presence-absence as a whole, instead of reporting the total number of individuals seen.

## Conclusions

These observed differences in biodiversity patterns serve as an important case study, highlighting the productivity and broad geographical reach of large, long-term, community science efforts. While the underlying causes of this data gap region remain a subject for future studies, the absence of such in the structured sampling alludes to a human-driven source of bias in community sampling efforts.

Although community science has been shown to be capable of generating large amounts of observations, the actual efficacy of community science-based reporting for this taxon appears to rely heavily on external factors pertaining to how people interact with nature. We expect that population density and accessibility may be a large predictive factor of community engagement, underscoring a need for further research into engagement patterns in community science efforts and potential biases that may arise from them. Ultimately, for studies interested in range and biodiversity, community science data could represent a thorough, crowd-sourced alternative to traditional data sources, especially in areas with prolific community initiatives.

Future efforts will be aimed at identifying and analyzing other sources of inaccuracies or biases within community science efforts. More particularly, future studies will focus on the effects of coloration and visibility of common odonates on community science reporting rates, as well as evaluating how community science efforts compare to historical museum collections for this region.

